# Robust Conditional Diffusion with Noisy Templates for Antibody Sequence–Structure Design

**DOI:** 10.64898/2026.06.18.733127

**Authors:** Peishuo Liu, Chenyang Yan, Mianzhi Pan, Xiao Liu, Shujian Huang, Zhen Wu, Jianbing Zhang

## Abstract

Antibodies specifically recognize antigens and play a central role in therapeutic discovery. Designing antibodies for a given antigen remains challenging because antigen–antibody complex data are limited, whereas the sequence and conformational spaces of complementarity-determining regions (CDRs) are large. Retrieved CDR templates from databases or candidate libraries can narrow the design space and improve controllability, but retrieval for novel antigens is often sparse and imperfect; treating retrieved templates as hard conditions can bias the denoising process and cause negative transfer. To address this problem, we propose Robust Conditional Diffusion with Noisy Templates for antibody sequence–structure design (NT-ABDiff), a joint diffusion framework that treats candidate CDR-only templates as optional and potentially unreliable conditions. NT-ABDiff uses reliability-aware template modulation to estimate the context-conditioned usefulness of each candidate and to adaptively reweight and fuse multiple templates during conditioning. We further train the model with mixed-quality and corrupted templates as conditional perturbation regularization, encouraging the denoiser to exploit informative templates while remaining stable when templates are uninformative. Experiments under controlled template shifts and a train-set retrieval evaluation show that NT-ABDiff improves CDR-H3 sequence recovery and structural accuracy over strong baselines, while retaining robustness to missing, mismatched, and corrupted templates. Under a stringent random-template CDR-H3 evaluation, NT-ABDiff improves amino-acid recovery (AAR) from 30.03% to 39.47% and reduces RMSD from 3.160 to 2.915 Å; with train-set retrieval candidates, it achieves 39.50% AAR and 2.76 Å RMSD. Code, processed splits, configuration files, and evaluation scripts are available at https://github.com/ShiDeng7rz/NT-ABDiff.

## 1 Introduction

ANTIBODIES are critical therapeutic proteins whose antigen binding is largely determined by specific interactions between their Complementarity-Determining Regions (CDRs) and target antigens [1], [2]. Consequently, antigenspecific CDR design can be formulated as a sequence– structure generation problem [3], [4], [5]. Computational approaches, particularly deep generative models, have become increasingly important for this task by reducing the search burden associated with time-consuming experimental screening [6], [7], [8], [9]. However, these models face substantial data and modeling challenges: the antibody sequence–structure space is vast, whereas available antigen– antibody complex data remain limited [10], [11]. Such data scarcity can hinder deep models from learning reliable mappings from antigen context to CDR sequence–structure configurations, leading to generated candidates with reduced structural accuracy or biological plausibility [12], [13].

To mitigate data sparsity and reduce the effective search space, one common strategy is to incorporate retrieved templates, such as homologous CDRs from databases, as inductive priors [8], [15]. In principle, templates can provide useful references and improve controllability. However, for novel antigens, retrieved candidates may be sparse, weakly matched, or inconsistent with the target antigen context, making rigid template conditioning unreliable. As shown in our nearest-neighbor analysis (Fig. 1), H1 and H2 often have closer template matches, whereas the critical CDR-H3 loop—which strongly influences binding specificity—shows substantially higher diversity and structural variability [14], [16].

**Fig. 1:**
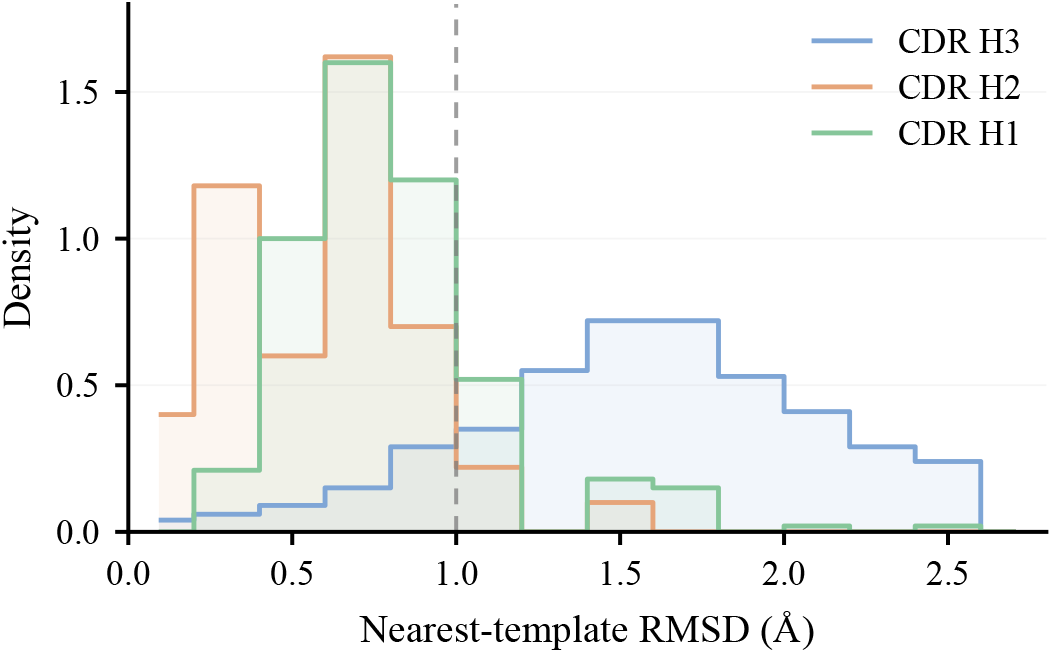
A nearest-neighbor view of CDR loop geometry suggests H1/H2 often have closer non-Ig neighbors than the highly diverse H3, making generic loop priors and template retrieval less reliable for H3 ( [14]). Consistently, the fraction of loops with nearest-template RMSD *<* 1Å is 0.82/0.93/0.14 for H1/H2/H3.

This creates a practical limitation for template-conditioned methods: when retrieved candidates are weakly matched, corrupted, or unrelated to the target context, rigid conditioning can introduce negative transfer by propagating misleading template signals into the denoising process. Conversely, discarding templates entirely removes potentially useful references and leaves generation to rely only on the antigen–framework context. A more robust conditioning mechanism should therefore estimate the context-conditioned usefulness of candidate templates, exploit informative candidates when available, and suppress unreliable candidates when they are corrupted or mismatched.

To address these challenges, we propose Robust Conditional Diffusion with Noisy Templates for Antibody Sequence–Structure Design (NT-ABDiff). The core idea is to treat candidate CDR templates as *optional, potentially unreliable* conditions (Fig. 2) and to model their contextdependent usefulness within a joint sequence–structure diffusion framework. Concretely, we encode multiple candidate CDR-only sequences into template features and fuse them with the antigen–framework context representation via multi-template attention. We further introduce a *reliability-aware template modulation* module, implemented as a template-wise learnable gate, that adaptively regulates the template-level contribution of each candidate during denoising. When a candidate template is compatible with the antigen–framework context, the gate retains or increases its contribution, allowing the template to provide an additional conditional signal for sequence–structure denoising. In contrast, when candidate templates are noisy, mismatched, or uninformative, their influence is suppressed. This reduces the risk of the model from forming rigid reliance on templates and keeps denoising primarily driven by the antigen-framework context. To expose the model to a controlled spectrum of template quality, we mix informative, corrupted, and uninformative template conditions during training. This encourages a conditional usage pattern in which templates are exploited when informative but are not required for stable generation. From this perspective, noisy templates serve as *conditional perturbations* that regularize training, and we empirically evaluate the resulting robustness under controlled template shifts and train-set retrieval candidates. NT-ABDiff is designed as a lightweight template-conditioning extension to a DiffAb-style sequence– structure diffusion backbone, rather than a replacement of the underlying diffusion formulation.

**Fig. 2:**
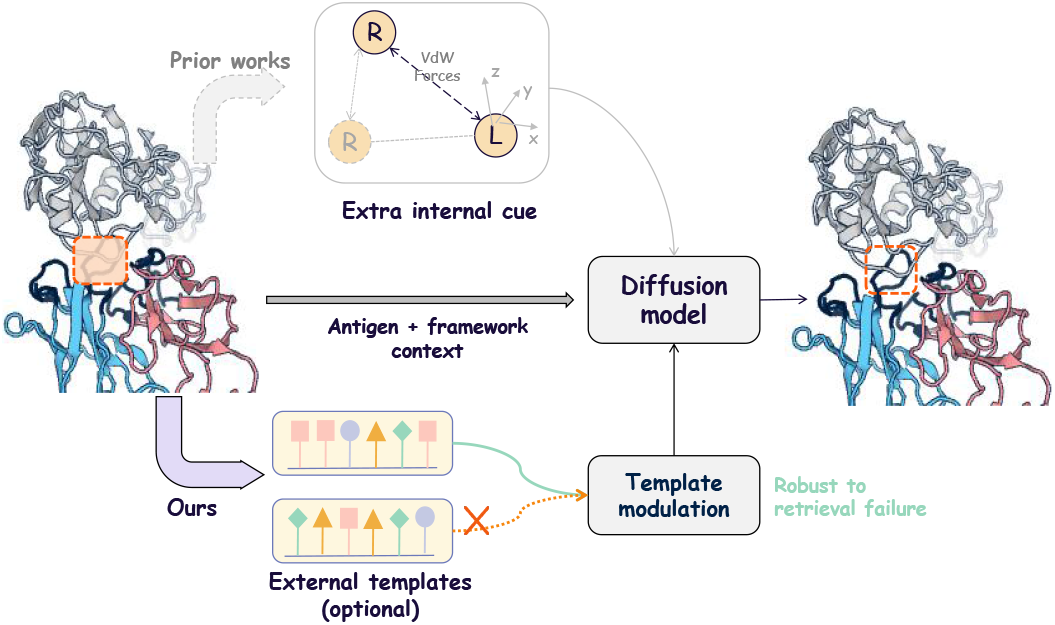
External optional template conditioning versus internal cue augmentation. Prior works strengthen diffusion with internal cues, such as physical or geometric features, whereas NT-ABDiff uses candidate CDR templates as optional weak conditions and suppresses uninformative candidates through reliability-aware template modulation.

The main contributions of this work are summarized as follows:

- **Robust conditioning with unreliable CDR templates**. We propose NT-ABDiff, a robust template-conditioning framework built on a joint sequence– structure diffusion backbone, which treats candidate CDR templates as *optional and potentially unreliable* conditions rather than rigid priors. We train and evaluate under controlled template-quality shifts and a train-set retrieval evaluation, showing improvements in CDR-H3 sequence recovery and structural accu-racy in unreliable-template settings. We also report a stringent CDR-H3-only evaluation protocol to reduce potential cross-CDR overlap effects and provide a more conservative assessment.
- **Reliability-weighted template modulation**. We introduce an adaptive conditioning module that estimates a context-conditioned usefulness score for each template and reweights candidates according to their compatibility with the antigen–framework context. This design reduces the risk of brittle dependence on template signals and allows denoising to rely primarily on the antigen–framework context when templates are uninformative.
- **Conditional-perturbation regularization**. We show that training with mixed-quality templates can be interpreted as a *conditional perturbation* regularizer that improves robustness of the denoising process. The resulting gains persist even when test-time templates are purely random, suggesting that the improvement is not solely attributable to direct template matching.

## 2 Related Work

### 2.1 Antibody Design and Antigen-Specific Co-design

Computational antibody design has long relied on template-based modeling and energy-function optimization, as exemplified by Rosetta-style pipelines [8], [17], [18], [19]. Recent learning-based methods can be broadly grouped into two categories. The first category focuses on sequence-centric design, including antibody language models [20], [21], repertoire-based priors, and structure-conditioned inverse folding [22], [23]. The second category focuses on antigenspecific sequence–structure co-design, where models operate on antibody–antigen complexes and jointly generate or predict CDR residue types and three-dimensional conformations [6], [24], [25], [26], [27]. Among these methods, DiffAb-style diffusion models provide an effective backbone for antigen-specific CDR generation by sampling complex-conditioned sequences and structures [6], [9], [28], [29].

Most existing co-design methods, however, rely primarily on the antigen–framework context and strengthen generation through internal conditioning signals, such as geometric features, physical cues, or sampling guidance. They do not directly address how external retrieved templates should be used when the retrieved candidates are weakly matched, corrupted, or uninformative. In practice, template retrieval for novel antigens may produce mixed-quality candidates, and rigidly injecting such candidates can introduce misleading signals into the denoising process. Our work builds on a DiffAb-style sequence–structure diffusion backbone, but focuses on the conditioning pathway: NT-ABDiff treats CDR templates as optional external conditions and uses a gated template-conditioning module to reduce brittle dependence on unreliable candidates.

### 2.2 Template Priors and Retrieval-Augmented Antibody Design

Template priors are widely used in protein and antibody structure modeling [30], but their utility depends on the availability and relevance of suitable templates. This issue is particularly important for CDR-H3, which is highly diverse and often has weaker nearest-neighbor similarity than H1 and H2 loops [14], [31]. Consistent with our nearest-neighbor analysis in Fig. 1, this property makes template transfer for H3 less reliable and motivates evaluating template-conditioned generation under degraded or weakly matched template conditions.

Retrieval-augmented diffusion has also been explored for antibody design. For example, RADAb uses retrieved structural priors to guide structure-informed antibody design [28]. This line of work is related to our use of external template information, but the input setting is different. Our task masks the target CDR sequence and structure and uses only the antigen–framework context together with optional CDR-only sequence templates. Therefore, methods that rely on target CDR structural information are not directly comparable under the same input constraints, although they provide relevant context for retrieval-augmented antibody design.

In this work, we study a complementary problem: how a sequence–structure diffusion model should use candidate CDR templates when their usefulness is variable. Rather than assuming that retrieved candidates are reliable priors, we treat them as noisy, optional auxiliary conditions. The learned gate is interpreted as a context-conditioned usefulness scorer, not as a calibrated uncertainty estimator. We train with a controlled spectrum of informative, corrupted, and uninformative templates, and further evaluate the learned conditioning behavior using train-set retrieval candidates. This design aims to exploit informative templates when available while allowing the denoising process to rely primarily on the antigen–framework context when template signals are unreliable.

## 3. Methods

### 3.1 Problem Formulation

We focus on joint sequence–structure generation of antibody CDRs, especially the highly variable CDR-H3. Each residue in an antibody–antigen complex is represented by three components for *i* = 1, …, *N* : residue identity *s*_*i*_*∈ A*, where denotes the 20 standard amino acids; three-dimensional coordinates **x**_*i*_ ∈ ℝ^3^; and spatial rotation **O**_*i*_ ∈*SO*(3).

Let the complex be decomposed into the antigen context and antibody regions. We denote the conditioning context as *C*:= (*C* _ag_, *C* _fr_), where *C* _ag_ is the antigen and *C*_fr_ is the antibody framework. The target CDR region is denoted by *C*_cdr_. To represent variable template quality, we optionally condition the model on a set of *K* candidate CDR-only sequence templates, 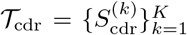, which may contain informative, corrupted, or uninformative candidates. Our goal is to model the conditional distribution of the target CDR state, *p*(*C* _cdr_, |*C T*_cdr_), i.e., to generate the CDR residue types and three-dimensional conformations conditioned on the antigen–framework context and the optional template set (Fig. 3).

**Fig. 3:**
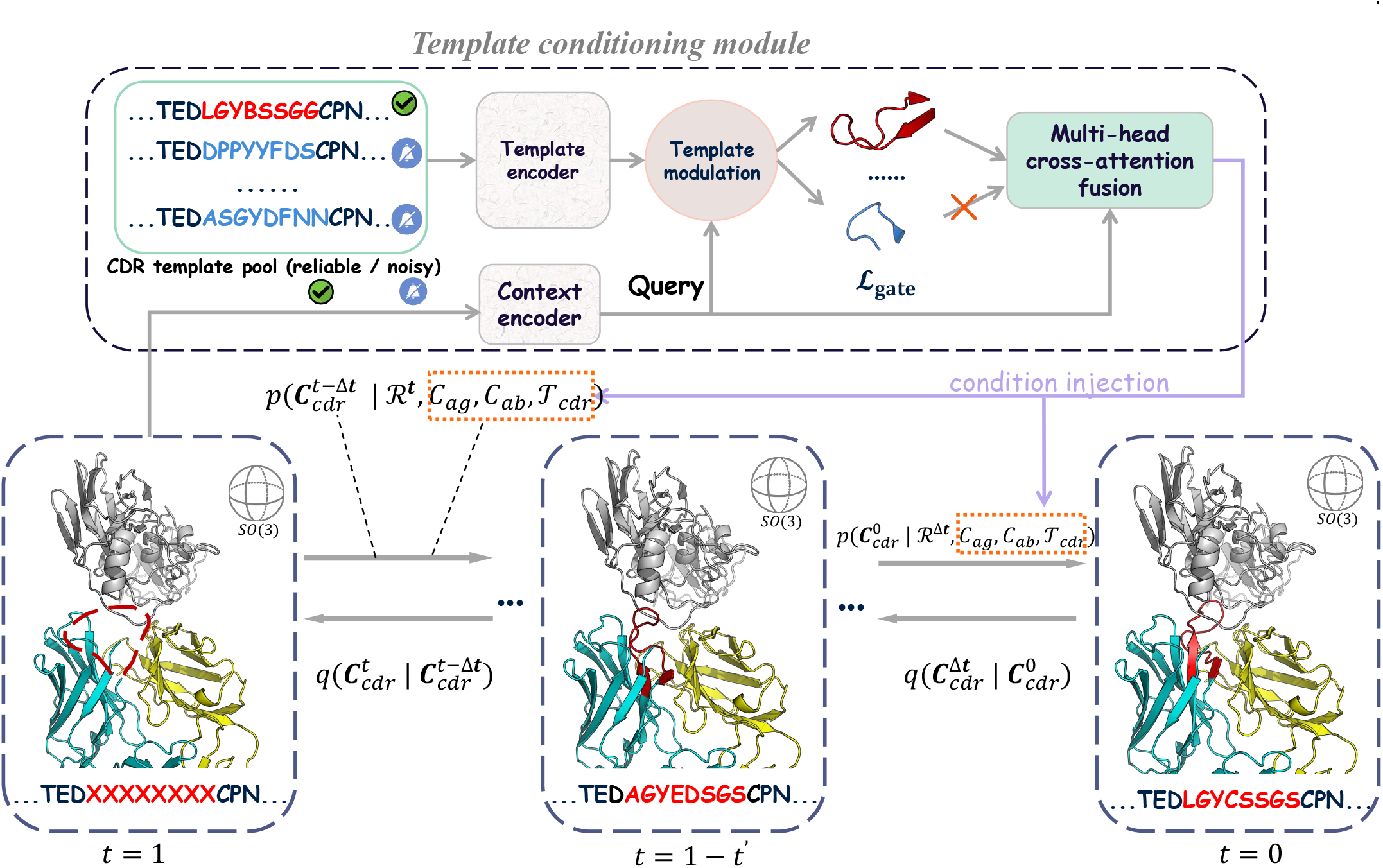
Overview of NT-ABDiff template conditioning. A pool of candidate CDR-only sequence templates is encoded and aligned to the full antibody length through padding, and a reliability-aware template modulation module, implemented as a template-wise gate, estimates the context-conditioned usefulness of each candidate. The modulated template features are injected through multi-template cross-attention followed by residual updates, enabling suppressible conditioning under mixed-quality template inputs.

### 3.2 Mixed-Quality Template Construction and Encoding

We follow DiffAb [6] as the diffusion backbone and augment its conditioning pathway with mixed-quality CDR-only templates to expose the model to variable template quality.

For each complex, we define a binary mask *M*_cdr_ ∈ {0, 1 }^*L*^ over the antibody sequence of length *L*, where *M*_cdr,*i*_ = 1 indicates a target CDR position, such as H3. We construct a candidate set of *K* CDR-only sequence templates that spans a controlled spectrum of template quality, including native, homologous-like, and uninformative candidates.

Specifically, the template set contains three components: (1) one native CDR sequence *S*^*⋆*^; (2) *⌈ρK⌉* homologous-like sequences, denoted by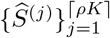, generated by applying random point mutations to *S*^*⋆*^; and (3) the remaining *K −⌈ρK⌉ −* 1 random sequences, denoted by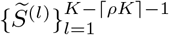,which serve as extreme uninformative controls. All templates are randomly shuffled to form the final batch:

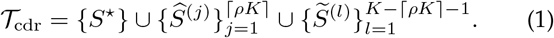

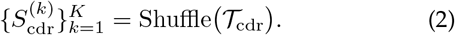

Each template 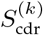 retains amino acid tokens only at CDR positions (where *M*_cdr,*i*_ = 1), with non-CDR positions filled by a special padding token `[PAD]`:

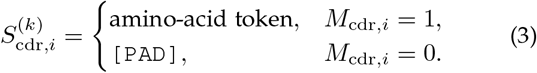

Each candidate template is then mapped to a continuous residue-aligned feature representation by a parameterized encoder ℰ_Θ_:

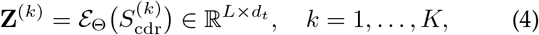

In practice, ℰ _Θ_ can be implemented using token embeddings, including `[PAD]`, optionally followed by lightweight contextual layers. This mixed-quality construction is used during training and is denoted as Trn-Mix. At inference, we evaluate controlled settings including Tst-Rand, where all *K* templates are random and no *S*^*⋆*^ is included, and Tst-Oracle, where the native template is included as an upper-bound condition. In addition, in Sec. 4, we evaluate train-set retrieval candidates produced by MMseqs2 to test whether the learned conditioning behavior transfers beyond synthetic template pools.

### 3.3 Reliability-Aware Template Modulation

At each diffusion step *t*, the backbone network encodes the antigen–framework complex into contextual residue representations 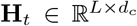 . To allow the model to determine *whether* and *to what extent* each candidate template should contribute under mixed-quality template inputs, we introduce a *template-wise learnable gating module*. For each candidate template, the gate outputs a scalar score *α*^(*k*)^ *∈* [0, 1], which is interpreted as a step-adaptive context-conditioned usefulness score rather than a calibrated uncertainty estimate:

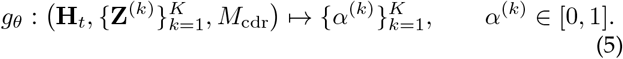

where 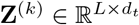 denotes the length-aligned template representation. Because *g*_*θ*_ conditions on **H**_*t*_, template con-tributions can be adapted across diffusion steps according to the current denoising state.

For each template *k*, the gating module first uses the residue-aligned contextual and template features and concatenates them along the feature dimension:

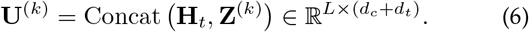

A lightweight interaction network *ϕ*_*θ*_(·) is then applied position-wise to extract interaction features:

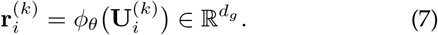

Because template sequences carry semantic information only at CDR residues, we aggregate interaction features using masked mean pooling over positions selected by *M*_cdr_:

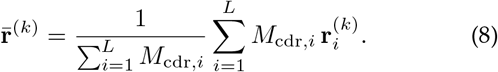

A scoring head *s*_*θ*_(·) then produces a template-level logit, followed by a sigmoid activation:

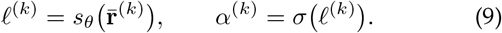

During training, we use a sigmoid-based straight-through estimator (STE) to obtain a hard gate 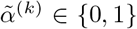 in the forward pass while allowing gradients to flow through the continuous relaxation *α*^(*k*)^ = *σ*(*l*^(*k*)^) [32]. At inference time, we apply a hard threshold 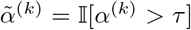. Unless otherwise specified, we use *τ* = 0.4 and analyze its sensitivity in Sec. 4. The gated template features are obtained by

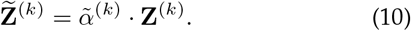

Overall, the gate makes template conditioning suppressible before cross-attention fusion: it retains or emphasizes candidates with high context-conditioned usefulness scores and suppresses candidates that appear mismatched or uninformative. Conditioning on **H**_*t*_ makes this modulation stepadaptive across diffusion and helps keep denoising driven by the antigen–framework context when template signals are unreliable.

### 3.4 Suppressible Multi-Template Conditioning

After gating, we incorporate the optional mixed-quality template features as a suppressible conditioning signal through a multi-template cross-attention fusion module. Let 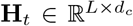 denote the contextual residue representations produced by the diffusion backbone at time step *t*, and let

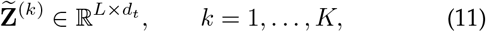

be the gated feature representation of the *k*-th template. We use **H**_*t*_ as the query and each template representation 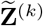 as the key and value to compute template-specific cross-attention responses:

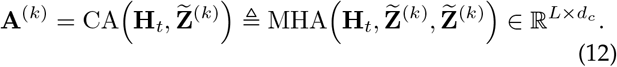

Here, CA(*·*) denotes a cross-attention operator whose output dimension matches that of the query, namely *d*_*c*_. Within multi-head attention, linear projections align the potentially different dimensions of contextual and template features.

Let 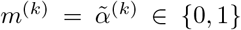 be the hard gate indicating whether template *k* is active. We define normalized mixture weights as

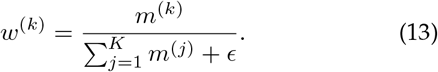

and fuse template-specific cross-attention outputs by a gated average:

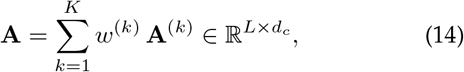

where *ϵ >* 0 is a small constant for numerical stability. If all templates are suppressed, then *w*^(*k*)^ = 0 for all *k* and **A** = **0**.

Finally, we inject the fused template message **A** into the contextual residue representations through a post-norm Transformer residual update:

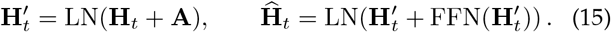

When all templates are suppressed, *m*^(*k*)^ = 0 for all *k*, so **A** = **0** and the template branch contributes no additional signal to the contextual representation. In our implementation, this template aggregation block is applied once per reverse step after the DiffAb context encoder and before concatenating the time embedding.

### 3.5 Joint Sequence–Structure Denoiser and Training Objective

We adopt DiffAb’s SE(3)-equivariant joint denoiser and reverse-process parameterization for residue types, coordinates, and orientations. Our method modifies only the conditional feature pathway: at each reverse step *t*, we first compute context-aware residue features **H**_*t*_ using DiffAb’s SE(3)-equivariant encoder, conditioned on the current (*R*_*t*_, *p*_*t*_) and the current sequence embedding, and then augment **H**_*t*_ with gated multi-template aggregation to ob-tain 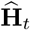 the diffusion process and diffusion loss remain unchanged. We then concatenate a time embedding *e*(*t*) with 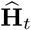and feed the resulting representation into three lightweight prediction heads for positional noise, rotational updates, and amino-acid posteriors.

For CDR positions indicated by the mask *M*_cdr_, we follow DiffAb and apply diffusion to orientations, coordinates, and residue types. At a sampled diffusion step *t ∼ U* ( { 1, …, *T*}), we minimize a weighted sum of three losses:

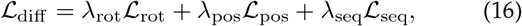

whereℒ _rot_ follows DiffAb and measures the discrepancy between predicted and target rotations on *SO*(3), ℒ _pos_ is the mean squared error (MSE) between the predicted and true positional noise, andℒ _seq_ is the KL divergence between the predicted amino-acid posterior and the forward-process posterior induced by the diffusion kernel.

To train the gate under the controlled mixed-quality template construction, we introduce an auxiliary binary supervision signal. Specifically, leveraging prior knowledge from the template construction process, we assign each candidate template *k* a binary label *y*^(*k*)^ ∈{0, 1}, where the native template and homologous-like point-mutated templates are labeled as 1, and randomly generated templates are labeled as 0. These labels are construction-specific proxies for template usefulness under our augmentation scheme, rather than calibrated uncertainty labels or general biological reliability annotations. In addition to producing the gating coefficient *α*^(*k*)^, the gating network also outputs an unnormalized logit *l*^(*k*)^ for each template. We supervise these logits using a binary cross-entropy on logits:

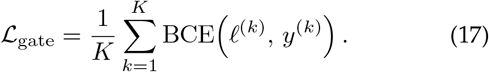

This auxiliary loss encourages the gating network to assign higher activation scores to informative templates, thereby forming a complement with the downstream generative task. Importantly, ℒ _gate_ serves as an auxiliary supervision on the gating logits and is optimized jointly withℒ _diff_ ; it does not change the diffusion formulation or the denoising objective of the DiffAb backbone. Here *y*^(*k*)^ is a synthetic training signal from our augmentation (native/homologous vs. random), and no labels are required at inference time. The final training objective is given by:

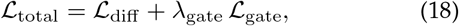

where *λ*_gate_ controls the relative contribution of the gate supervision term. By optimizingℒ _total_, the model learns sequence–structure denoising together with a suppressible template-conditioning policy for mixed-quality template inputs.

## 4 Experiments

We evaluate our model under a joint H1/H2/H3 sequence– structure co-design setting. In addition to the joint H1/H2/H3 evaluation, we report a more stringent CDRH3-only protocol to reduce potential cross-CDR overlap effects, since the split is clustered by H3 identity. This protocol offers a more conservative assessment for the highly variable CDR-H3 design setting (Table 2). We further include ablations and analyses to isolate the effects of mixed-quality template training and template-wise gating, and we evaluate train-set retrieval candidates to test whether the learned conditioning behavior transfers beyond synthetic template pools.

### 4.1 Experimental Setup

#### Dataset

We construct the dataset from SAbDab [10], removing complexes with resolution worse than 4.0 *Å* and antibodies binding non-protein antigens. Antibody residues are renumbered using ANARCI with the Chothia scheme [33], [34]. Following prior work, we cluster antibodies at 50% CDR-H3 sequence identity using MMseqs2 and split clusters into training and test sets to reduce homologous overlap across splits; test complexes with homologous training counterparts are removed. We additionally hold out 19 training complexes for validation. Unless otherwise specified, all experiments use the same split and preprocessing.

#### Evaluation protocol

For each test complex, we mask the target CDR region in both sequence and structure and require the model to jointly generate its amino-acid sequence and three-dimensional backbone, conditioned on the remaining framework and antigen context. We fix the CDR length to the native length for controlled comparison and generate candidates by sampling from the diffusion process. We sample 100 candidates per target CDR and apply a standard refinement pipeline to all generated structures: we first perform geometry relaxation with OpenMM [35] and then run Rosetta all-atom refinement and scoring [36]. For energy-based metrics, we compute binding/interface energy on the refined complexes to ensure a fair comparison across methods.

#### Baselines

We compare against representative approaches spanning diffusion-based co-design and classical energy-based design: DiffAb [6] (diffusion-based sequence– structure generation), DIFFFORCE [37] (force-guided diffusion sampling), RAbD [8] (Rosetta energy-based antibody design), and AbMEGD [9] (equivariant graph diffusion for complex binding). All diffusion baselines use the same candidate budget (100 samples) and the same refinement/scoring pipeline for a fair comparison.

#### Metrics

We report three metrics. **AAR** measures sequence recovery on the target CDR residues (amino-acid recovery rate) [8]. **RMSD** measures structural accuracy as C_*α*_ deviation after framework alignment [38]. **IMPROVE%** reports the fraction of designed CDRs whose refined complex achieves lower binding/interface energy than the native CDR under the same scoring protocol [36]. Main tables report benchmark means; seed-level variability for NT-ABDiff is reported in the supplementary material.

#### Implementation details

Unless otherwise specified, training and inference details are provided in the supplementary material. Briefly, we use *T* = 100 diffusion steps,optimize with Adam using a batch size of 16, use *K* = 8 candidate templates, and set the inference-time hard gate threshold to *τ* = 0.4.

Because the diffusion backbone, candidate budget, and refinement/scoring pipeline are kept fixed where applicable, the experiments mainly isolate the effect of mixed-quality template training and gated suppressible conditioning.

### 4.2 Sequence–Structure Co-design

To assess performance under uninformative templates and estimate an upper-bound condition, we evaluate NT-ABDiff under two settings: (1) **Rand**, where all templates are random candidates and the setting is used for fair comparison, and (2) **Oracle**, where a native template is included among the *K* candidates and the result is reported only as an upperbound condition.

Under the **Rand** setting, NT-ABDiff obtains the best fair-setting AAR on CDR-H1 (69.63) and CDR-H3 (38.96), with the H3 gain over AbMEGD being marginal (38.96 vs. 38.93), while remaining comparable on CDR-H2 (55.34 vs. 55.40 for AbMEGD). For structural accuracy, NT-ABDiff achieves the best fair-setting RMSD on CDR-H1 (1.141) and the second-best RMSD on CDR-H3 (3.065), where it improves over DiffAb (3.377), DIFFFORCE (3.612), and AbMEGD (3.419), although RAbD obtains a lower H3 RMSD (2.900). These results suggest that the suppressible template pathway improves sequence recovery without degrading CDRH3 structural accuracy relative to diffusion-based baselines. For IMPROVE%, NT-ABDiff is not the strongest method, which is expected because it does not explicitly optimize the Rosetta energy objective; nevertheless, on CDR-H3 it outperforms DiffAb (25.52 vs. 22.94) and AbMEGD (25.52 vs. 23.74), while remaining below DIFFFORCE (30.22).

Under the **Oracle** setting, including the native template further improves H1 and H3 in both AAR and RMSD (e.g., H3 RMSD: 2.994 vs. 3.065 in Rand), suggesting that the gated multi-template fusion can exploit informative templates when they are available. On CDR-H2, Oracle improves RMSD (1.122 vs. 1.188) and IMPROVE% (35.60 vs. 31.11), but does not improve AAR (54.69 vs. 55.34), suggesting that H2 may be more constrained by the framework– antigen context and may benefit less from sequence-level template information.

Taken together, these results indicate that NT-ABDiff improves key sequence-recovery metrics under uninformative template inputs, while retaining the ability to benefit from informative templates in the oracle upper-bound setting.

### 4.3 CDR-H3-only Evaluation and Ablation

Under a stricter protocol, we train and evaluate *only* on CDR-H3 to reduce potential cross-CDR overlap effects when clustering is performed solely by CDR-H3 identity. In the joint H1/H2/H3 setting, similar or overlapping H1/H2 contexts may remain across splits even when H3 identity is controlled, which can make the evaluation less conservative. Under the H3-only protocol, DiffAb also improves relative to the joint setting (AAR/RMSD: 30.03/3.160 vs. 27.02/3.377 in Table 1), suggesting that the focused H3-only setting provides a different and more conservative testbed for assessing CDR-H3 design.

**TABLE 1:** CDR sequence–structure co-design results under the joint H1/H2/H3 protocol. Best and second-best results among fair, non-oracle settings are shown in **bold** and <u>underlined</u>, respectively. Values are benchmark means. Five-seed variability of NT-ABDiff is reported in the supplementary material. *†*denotes an oracle-template setting that includes the native template as an upper-bound condition and is not used for ranking.

| Method | AAR (% , ↑) | | | RMSD ($\text{\AA}$ , ↓) | | | IMPROVE% (% , ↑) | | |
| --- | --- | --- | --- | --- | --- | --- | --- | --- | --- |
|  | H1 | H2 | H3 | H1 | H2 | H3 | H1 | H2 | H3 |
| RABD | 22.85 | 22.85 | 22.14 | 2.261 | 1.641 | **2.900** | 43.88 | **53.50** | 23.25 |
| DiffAb | 66.99 | 49.18 | 27.02 | 1.178 | **1.067** | 3.377 | 47.42 | 32.23 | 22.94 |
| DIFFFORCE | 60.78 | 53.51 | 29.52 | 1.561 | 1.401 | 3.612 | <u>49.45</u> | <u>36.81</u> | **30.22** |
| AbMEGD | 66.96 | **55.40** | <u>38.93</u> | <u>1.169</u> | <u>1.130</u> | 3.419 | **55.11** | 24.05 | 23.74 |
| **NT-ABDiff (Rand)** | **69.63** | <u>55.34</u> | **38.96** | **1.141** | 1.188 | <u>3.065</u> | 49.00 | 31.11 | <u>25.52</u> |
| *NT-ABDiff (Oracle)<sup>†</sup>* | 70.39 | 54.69 | 39.42 | 1.121 | 1.122 | 2.994 | 50.63 | 35.60 | 28.10 |

In this fair setting, NT-ABDiff with random test-time templates (**Tst-Rand**) improves over DiffAb (AAR: 39.47 vs. 30.03; RMSD: 2.915 vs. 3.160), indicating that the model does not require informative test-time templates to improve CDR-H3 sequence–structure generation. This result supports the intended conditioning behavior: templates should be useful when informative but should not be required when the provided candidates are uninformative. Moreover, when the native template is included (**Tst-Oracle**, upper bound), performance further improves (AAR/RMSD: 41.70/2.871), suggesting that the gated fusion module can exploit informative template signals when they are present. We report this setting only as an upper bound and do not treat it as directly comparable to fair baselines.

In the training ablations, removing the gating module substantially degrades performance (**Trn-Mix w/o Gate**: 29.87/3.194), bringing it close to DiffAb, indicating that the gate is important for suppressing uninformative template signals during conditional injection. Training with an overly idealized template source (**Trn-TrueOnly**^*\**^) yields the worst results (20.29/3.321), consistent with a train–test mismatch: seeing only perfect templates during training may encourage brittle reliance on template signals, which generalizes poorly when templates are imperfect at test time. Finally, **Trn-RandOnly** yields a small gain over DiffAb without gating (31.08/3.110), suggesting that random templates can act as a regularizing conditional perturbation during training. Adding gating brings little additional benefit in this case (31.56/3.107), consistent with the gate converging to near-constant suppression when all candidates are uninformative.

**TABLE 2:** Ablation on gated template conditioning under the CDR-H3-only protocol. Tst-Rand uses purely random templates and serves as the fair test-time setting; Tst-Oracle includes one native template and is reported only as an upperbound condition. Trn-Mix denotes mixed-quality template training with native, homologous-like, and random templates; Trn-RandOnly uses random-only templates; and Trn-TrueOnly^*\**^ uses only the native template. Best results among fair, non-oracle settings are shown in **bold**.

| Method | Train config | Test config | AAR (% , ↑) | RMSD ($\text{\AA}$ , ↓) |
| --- | --- | --- | --- | --- |
| DiffAb (baseline) | – | – | 30.03 | 3.160 |
| *Test-time template availability* |  |  |  |  |
| NT-ABDiff | Trn-Mix + Gate | Tst-Rand | **39.47** | **2.915** |
| NT-ABDiff<sup>†</sup> | Trn-Mix + Gate | Tst-Oracle | 41.70 | 2.871 |
| *Training-time ablations* |  |  |  |  |
| Trn-TrueOnly* | Trn-TrueOnly | Tst-Oracle | 20.29 | 3.321 |
| Trn-Mix w/o Gate | Trn-Mix (w/o Gate) | Tst-Rand | 29.87 | 3.194 |
| Trn-RandOnly w/o Gate | Trn-RandOnly (w/o Gate) | Tst-Rand | 31.08 | 3.110 |
| Trn-RandOnly + Gate | Trn-RandOnly + Gate | Tst-Rand | 31.56 | 3.107 |

Overall, the H3-only protocol provides a conservative assessment: NT-ABDiff improves CDR-H3 generation under uninformative template inputs, benefits from informative templates in the oracle upper-bound setting, and relies on gating to make template conditioning suppressible.

### 4.4 Train-Set Retrieval Evaluation

The controlled Tst-Rand and Tst-Oracle settings in Table 2 isolate the behavior of the conditioning module under uninformative and upper-bound template inputs. However, they do not fully address whether the learned conditioning policy transfers to templates obtained by sequence-based retrieval. To evaluate this point, we further construct a trainset retrieval setting using MMseqs2 candidates. For each test case, we use the heavy-chain non-H3 context as the query and retrieve candidate templates only from the training set. Retrieved candidates are filtered by sequence identity, matched H3 length, deduplication, and top-ranked selection. This protocol avoids using the test CDR-H3 sequence or structure as a query, while providing a retrieval-based template source beyond the synthetic template pools used in the controlled ablations.

Table 3 reports the resulting CDR-H3-only performance. Under the matched Trn-RealBank/Tst-RealBank setting, enabling the gate improves AAR from 30.08 to 31.76 and reduces RMSD from 3.18 to 3.13. This modest but consistent improvement suggests that the learned gate provides benefit beyond static retrieval filtering alone. More importantly, the model trained with mixed-quality synthetic templates (**Trn-Mix**) transfers directly to real-bank retrieval candidates and achieves 39.50 AAR and 2.76 Å RMSD on Tst-RealBank, outperforming the RealBank-trained variants. This result suggests that mixed-quality template training does not merely overfit to synthetic random/oracle pools; instead, it learns a suppressible conditioning policy that remains effective when candidate templates are obtained from trainset retrieval.

**TABLE 3:**
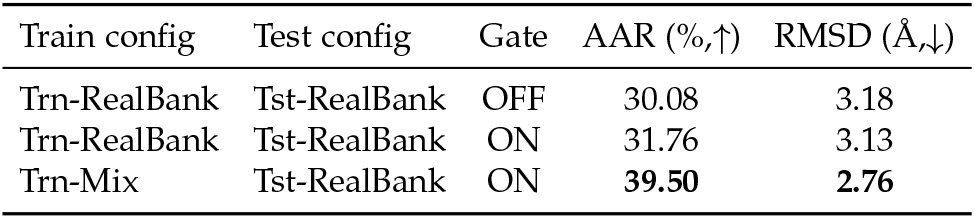
Train-set retrieval evaluation with MMseqs2 candidates under the CDR-H3-only protocol. Candidate templates are retrieved from the training set using the heavychain non-H3 context as query and filtered by pident ≥ 80, same H3 length, deduplication, and top-4 ranking. This setting evaluates transfer beyond synthetic template pools, but is not a full end-to-end retrieval system over an external database.

| Train config | Test config | Gate | AAR (% , ↑) | RMSD ($\text{\AA}$ , ↓) |
| --- | --- | --- | --- | --- |
| Trn-RealBank | Tst-RealBank | OFF | 30.08 | 3.18 |
| Trn-RealBank | Tst-RealBank | ON | 31.76 | 3.13 |
| Trn-Mix | Tst-RealBank | ON | **39.50** | **2.76** |

This experiment should be interpreted as a train-set retrieval evaluation rather than a full end-to-end retrieval benchmark over an external antibody database. Nevertheless, it addresses the main concern that the proposed conditioning mechanism only works under artificial template construction: the same Trn-Mix model remains effective when evaluated with MMseqs2-retrieved candidates, and the gate continues to provide gains after retrieval-side filtering.

### 4.5 Gate Behavior under Mixed and Random Template Pools

We analyze additional statistics of the gating module *g*_*θ*_ (Fig. 4 and Table 4) under two controlled template regimes: (i) **mixed pools** used for diagnostic analysis, where construction-positive templates, including native and homologous-like point-mutated candidates (*y*=1), are mixed with random candidates (*y*=0); and (ii) **randomonly** templates (Tst-Rand), which serve as an extreme uninformative-template control in the fair protocol.

**TABLE 4:** Gate statistics for Fig. 4 with *τ* = 0.4. The gate score *α* is interpreted as a construction-specific template-usefulness score, not as a calibrated uncertainty probability.

| Setting | Mean | Median | $\Pr(\alpha > \tau)$ |
| --- | --- | --- | --- |
| Mixed: $y = 1$ | 0.331 | 0.407 | 58.33% |
| Mixed: $y = 0$ | 0.0437 | 0.00921 | 2.9% |
| Rand-only, all | 0.071 | 0.015 | 3.9% |
| Rand-only, top-1 | 0.323 | – | 32.8% |

**Fig. 4:**
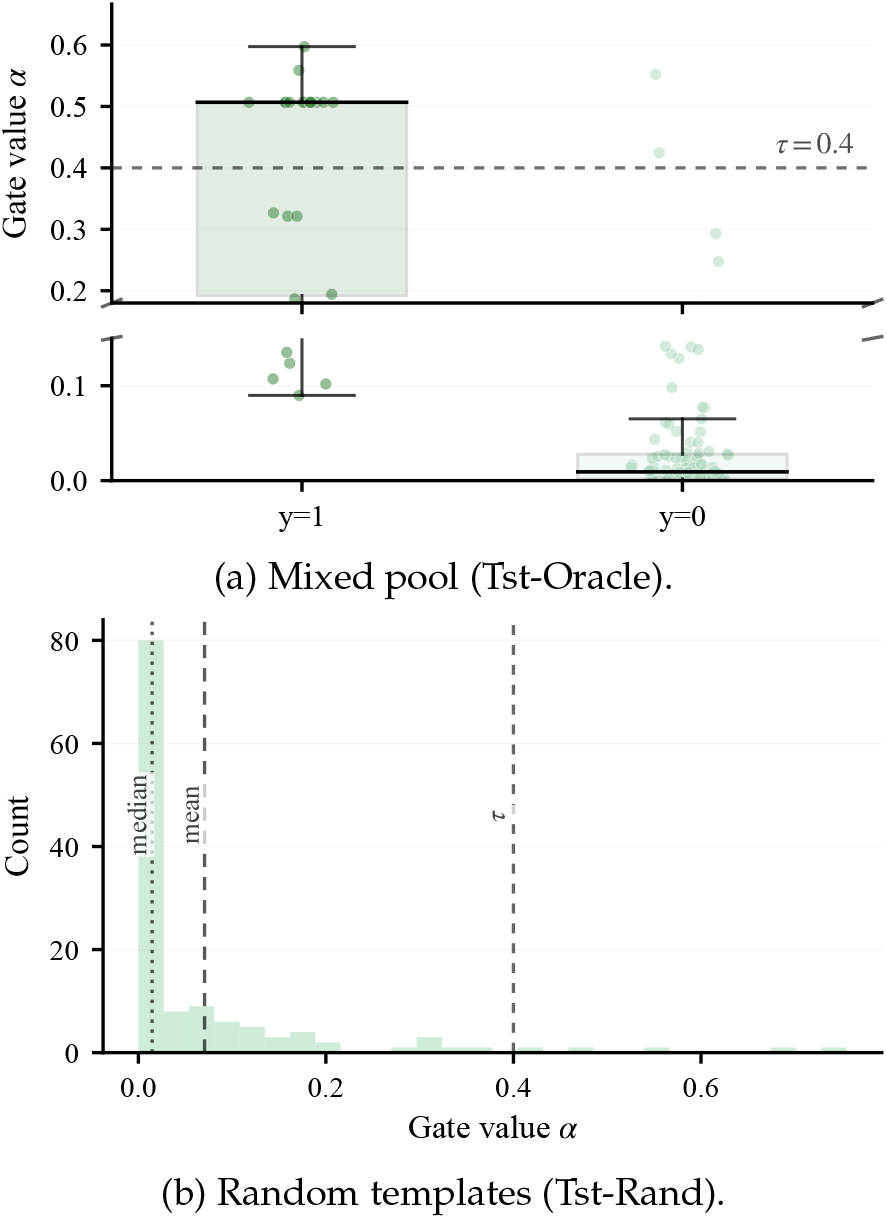
Gating behavior under noisy templates. **(a)** In a mixed pool containing one native/homologous candidate, the gate assigns larger coefficients *α* to templates labeled *y*=1 than to random templates (*y*=0). **(b)** Under random-only templates, *α* concentrates near zero (with mean/median marked), and the threshold *τ* =0.4 rejects most candidates. Together, these results suggest context-dependent template selection rather than hard filtering.

#### Mixed pools

The sigmoid gate scores *α* = *σ*(*l*) assign substantially higher values to *y* = 1 candidates than to *y* = 0 candidates under the mixed-pool construction: the ratio of mean scores is approximately 7.5 *×*(0.331 vs. 0.0437), and the fractions above the threshold *τ* = 0.4 differ markedly (58.33% vs. 2.9%; Table 4). In the analyzed mixed-pool subset, the top-1 gated candidate corresponds to a construction-positive candidate in 11/12 cases, suggesting that *g*_*θ*_ usually ranks informative candidates above random controls when such candidates are present in this diagnostic setting.

#### Random-only gate activations

Under Tst-Rand, gate scores are concentrated near zero (mean 0.071, median 0.015), with a small tail of higher-scoring candidates. Although only 3.9% of random *candidates* exceed *τ* = 0.4, evaluating *K* candidates per sample increases the per-sample maximum;consequently, 32.8% of *samples* contain at least one candidate with top-1 *α > τ* . This behavior is consistent with a context-conditioned scoring mechanism rather than a hard detector: even among random controls, a small subset can receive nonzero scores due to accidental compatibility with the current antigen–framework representation.

#### Why occasional activations do not dominate denoising

Occasional activations are compatible with the suppressible conditioning design: the DiffAb-style backbone remains the primary denoising pathway, while template signals are gated before cross-attention and injected through residual updates. Empirically, NT-ABDiff remains effective under Tst-Rand (Table 2), whereas removing the gating module degrades performance, indicating that gating is important for suppressing uninformative template signals during conditional injection.

### 4.6 Sensitivity to the Number of Candidate CDR Templates

To examine sensitivity to the number of candidate CDR templates, we vary *K* while keeping the other training conditions fixed, including the template mixing ratio, CDR-H3only training protocol, and 90k-iteration training budget. Each setting is repeated with three random seeds to reduce the influence of stochastic variation and to support selection of the default template number.

#### Effect of template number

As shown in Fig. 5(a) and Fig. 5(b), using candidate templates improves RMSD and AAR over the DiffAb baseline (RMSD = 3.160, AAR = 30.03) for the tested *K* values. Among the tested settings, *K* = 8 gives the best overall performance, which motivates using *K* = 8 as the default in the main experiments.

**Fig. 5:**
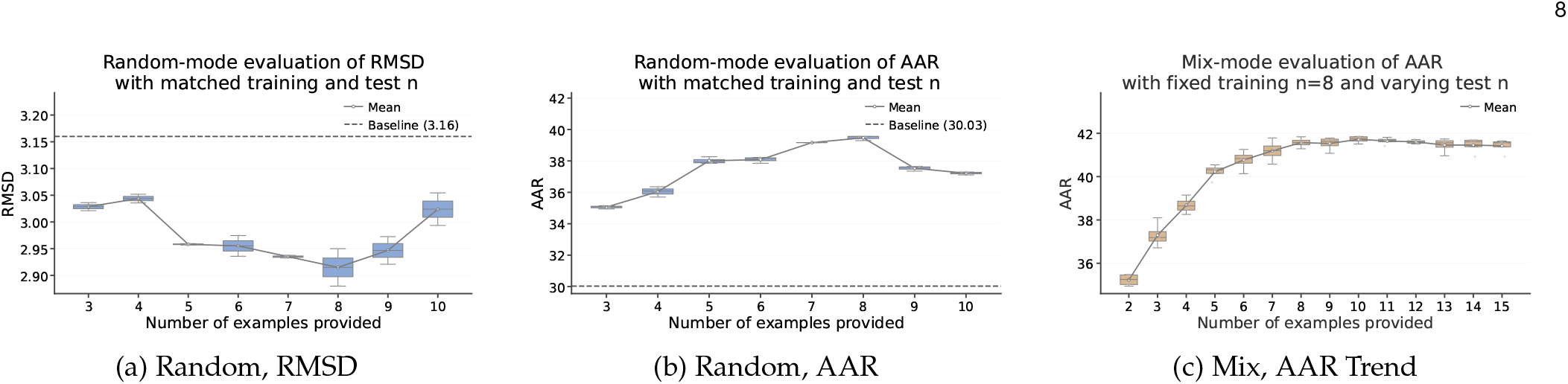
Sensitivity to the number of candidate CDR templates. Subplots (a) and (b) report RMSD and AAR when varying the number of candidate templates *K* under the CDR-H3-only protocol. Subplot (c) shows the test-time trend when the model is trained with *K* = 8 and evaluated with different numbers of candidate templates.

#### Test-time saturation with K = 8 training

When the model is trained with *K* = 8, Fig. 5(c) shows that performance improves as the number of test-time candidate templates increases. The gain is more pronounced when increasing from very few templates, and the curve gradually saturates around *K* = 8, suggesting diminishing returns from providing additional candidate templates within the tested range.

## 5 Conclusion

We presented NT-ABDiff, a lightweight template-conditioning extension to a DiffAb-style sequence–structure diffusion backbone for antigen-specific CDR design under potentially unreliable CDR template inputs. NT-ABDiff treats CDR-only templates as optional conditions, fuses multiple candidates through multi-template attention, and modulates their influence with a template-wise gate during denoising. Training with mixed-quality templates improves performance under controlled uninformative-template settings and transfers to train-set retrieval candidates. Across the joint benchmark and the stricter CDR-H3-only protocol, NT-ABDiff improves key sequence-recovery and structural metrics under uninformative template inputs, while the oracle setting shows that the model can further benefit from informative templates when available. Gate analyses show that the learned scores separate construction-positive templates from random controls in diagnostic mixed pools and suppress most random candidates under the random-only setting, supporting the interpretation of the gate as a construction-specific template-usefulness scorer rather than a calibrated uncertainty estimator.

### Limitations and ethical considerations

The evaluation is limited by the size and bias of available antigen–antibody complex datasets, which are enriched for well-studied targets; performance may degrade for rare epitopes or template sources that differ substantially from the evaluated settings. Our controlled template-shift settings use random, mixed-quality, and corrupted templates, and the retrieval experiment uses train-set retrieval candidates rather than a full end-to-end retriever over a large disjoint external database. The current model also does not explicitly optimize developability properties, such as stability, immunogenicity, or manufacturability, and does not guarantee binding improvement; generated candidates therefore require downstream filtering and experimental validation. This work evaluates the conditioning module on a DiffAb-style backbone; although the module is lightweight and modular, transfer to other antibody generative backbones remains to be validated.

Because biomolecular generative models may raise dual-use concerns, model release and use should follow institutional, legal, and community safety policies.

