## Supplementary Material for "Robust Conditional Diffusion with Noisy Templates for Antibody Sequence–Structure Design"

### S1 SAbDab Data and Preprocessing

#### S1.1 SAbDab as the Data Source

SAbDab is a curated antibody-structure database that provides antibody–antigen complex annotations, including heavy–light chain pairing, antigen chain information, and structure-level metadata. These annotations are useful for our setting because each paired antibody–antigen complex can be treated as one training or evaluation instance, with the antigen and antibody framework forming the conditioning context and the CDR region forming the generation target.

Beyond structural coordinates, SAbDab provides metadata that supports reproducible dataset construction. We fix the SAbDab snapshot downloaded in April 2025 and record the PDB IDs used in the experiments. We then use experimental and annotation fields to apply the same filtering protocol as in the main paper: removing complexes with resolution worse than $4.0\,\AA$, excluding antibodies binding non-protein antigens, and removing entries with unresolved required chains or CDR residues. Antibody residues are renumbered using ANARCI with the Chothia numbering scheme, which allows consistent extraction of CDR-H1/H2/H3 regions and construction of residue masks across heterogeneous complexes.

#### S1.2 Dataset Bias and Split Control

As with most structure-derived resources, SAbDab reflects historical and experimental biases: some antigen families and antibody classes are more heavily represented than others, while many clinically relevant targets have limited structural coverage. This motivates careful split construction and conservative evaluation. Following the main paper, we cluster antibodies at 50% CDR-H3 sequence identity using MMseqs2 and split clusters into training and test sets to reduce homologous overlap across splits. We also remove test complexes with homologous training counterparts and hold out 19 training complexes for validation.

This split strategy does not eliminate all possible similarities across antibody contexts, especially outside CDR-H3. Therefore, the main paper additionally reports a CDR-H3-only protocol to reduce potential cross-CDR overlap effects and provide a more conservative assessment for the highly variable CDR-H3 design task. The mixed-quality template construction should be interpreted as a controlled template perturbation protocol for studying suppressible conditioning, rather than a complete simulation of all possible retrieval-error modes in external antibody databases.

### S2 Details of Diffusion Processes

This appendix summarizes the joint sequence–structure diffusion formulation of DiffAb, which we use as our generative backbone. It is included only for completeness and to fix notation and does not introduce new technical contributions.

#### S2.1 Setup and Notation

We consider an antibody-antigen complex with a fixed context $\mathcal{C}$ (antigen structure and antibody framework) and a set of CDR residues $\mathcal{C}_{\mathrm{cdr}}$ to be generated. Each residue $j \in \mathcal{C}_{\mathrm{cdr}}$ is represented by a triplet $(s_j, \mathbf{x}_j, \mathbf{O}_j)$ where $s_j \in \{1, \ldots, 20\}$ is the amino acid type, $\mathbf{x}_j \in \mathbb{R}^3$ is the $\mathrm{C}\alpha$ coordinate, and $\mathbf{O}_j \in SO(3)$ is the residue orientation. Let $\mathbf{R}^t = \{(s_j^t, \mathbf{x}_j^t, \mathbf{O}_j^t)\}_{j \in \mathcal{C}_{\mathrm{cdr}}}$ denote the CDR state at the diffusion step $t \in \{0, \ldots, T\}$, where $t = 0$ is the data state and $t = T$ is close to the factorized noise prior.

**Templates as optional conditions.** In our method, we additionally condition the denoisers on a set of candidate CDR-only templates $\mathcal{T} = \{S_k\}_{k=1}^{K}$. For notational compactness, we absorb templates into the condition and write $\tilde{\mathcal{C}} := (\mathcal{C}, \mathcal{T})$; throughout this appendix we use $\mathcal{C}$

*P. Liu, C. Yan, M. Pan, X. Liu, S. Huang, Z. Wu, and J. Zhang are with the National Key Laboratory for Novel Software Technology, Nanjing University, Nanjing, Jiangsu, China.*
*P. Liu and C. Yan contributed equally to this work.*
†*Corresponding author: Jianbing Zhang.*

as a shorthand for $\tilde{\mathcal{C}}$ when no ambiguity arises. **Density shorthand.** To save space, we write $\mathcal{N}(\mu, \Sigma)$ as shorthand for the Gaussian density $\mathcal{N}(x \mid \mu, \Sigma)$ when the random variable is clear from context. Similarly, following DiffAb, $\mathrm{IGSO}(3)(\mathbf{M}, \sigma^2)$ denotes the isotropic Gaussian density on $SO(3)$ with mean rotation $\mathbf{M} \in SO(3)$ and scalar variance $\sigma^2$ (again omitting the explicit "|" form). We use the same shorthand throughout this appendix.

### S2.2 Forward Diffusion Processes

Following DiffAb, we define three independent forward (corruption) processes over the CDR variables $(s, \mathbf{x}, \mathbf{O})$—residue type, C$\alpha$ position, and orientation—with time-dependent noise schedules $\{\beta_t^\star\}_{t=1}^T$ for $\star \in \{\text{type}, \text{pos}, \text{ori}\}$. As in our main method, the forward diffusion is applied only to CDR positions indicated by $M_{\text{cdr}}$, while templates enter the model purely through conditioning. Define cumulative products

$$\bar{\alpha}_t^\star = \prod_{\tau=1}^{t} (1 - \beta_\tau^\star), \quad \star \in \{\text{type}, \text{pos}, \text{ori}\}.$$

**Multinomial diffusion for amino acid types.** For each residue $j$, the forward kernel replaces the type with a uniform amino acid draw with probability $\beta_t^{\text{type}}$ (and keeps it unchanged otherwise):

$$q(s_j^t \mid s_j^{t-1}) = \text{Multinomial}\Big((1 - \beta_t^{\text{type}})\,\text{onehot}(s_j^{t-1}) + \beta_t^{\text{type}}\,\tfrac{1}{20}\mathbf{1}\Big). \tag{S1}$$

where $\text{onehot}(\cdot) \in \mathbb{R}^{20}$ and $\mathbf{1}$ is the all-ones vector. By composing the kernels over steps, the closed-form marginal becomes

$$q(s_j^t \mid s_j^0) = \text{Multinomial}\Big(\bar{\alpha}_t^{\text{type}}\,\text{onehot}(s_j^0) + (1 - \bar{\alpha}_t^{\text{type}})\,\tfrac{1}{20}\mathbf{1}\Big). \tag{S2}$$

**Gaussian diffusion for C$\alpha$ coordinates.** The coordinate forward kernel is an isotropic Gaussian:

$$q(\mathbf{x}_j^t \mid \mathbf{x}_j^{t-1}) = \mathcal{N}\Big(\sqrt{1 - \beta_t^{\text{pos}}}\,\mathbf{x}_j^{t-1},\ \beta_t^{\text{pos}}\mathbf{I}\Big), \tag{S3}$$

where $\mathbf{I}$ is the $3 \times 3$ identity matrix and the closed-form marginal is

$$q(\mathbf{x}_j^t \mid \mathbf{x}_j^0) = \mathcal{N}\Big(\sqrt{\bar{\alpha}_t^{\text{pos}}}\,\mathbf{x}_j^0,\ (1 - \bar{\alpha}_t^{\text{pos}})\mathbf{I}\Big). \tag{S4}$$

Equivalently, with $\boldsymbol{\epsilon}_j \sim \mathcal{N}(\mathbf{0}, \mathbf{I})$,

$$\mathbf{x}_j^t = \sqrt{\bar{\alpha}_t^{\text{pos}}}\,\mathbf{x}_j^0 + \sqrt{1 - \bar{\alpha}_t^{\text{pos}}}\,\boldsymbol{\epsilon}_j. \tag{S5}$$

**Diffusion on $SO(3)$ for orientations.** For orientations, DiffAb specifies a closed-form forward perturbation on $SO(3)$ via an isotropic Gaussian:

$$q(\mathbf{O}_j^t \mid \mathbf{O}_j^0) = \mathrm{IGSO}(3)\Big(\text{ScaleRot}(\sqrt{\bar{\alpha}_t^{\text{ori}}}, \mathbf{O}_j^0),\ 1 - \bar{\alpha}_t^{\text{ori}}\Big), \tag{S6}$$

where $\text{ScaleRot}(\cdot, \cdot)$ scales the rotation angle while keeping the rotation axis fixed.

### S2.3 Reverse (Generative) Processes

The reverse (generative) kernels are parameterized by neural networks conditioned on the noisy CDR state $\mathbf{R}^t$, the diffusion step $t$ (via a time embedding), and the conditioning context $\tilde{\mathcal{C}} := (\mathcal{C}, \mathcal{T})$; for brevity we write $\mathcal{C}$ for $\tilde{\mathcal{C}}$ when no ambiguity arises. In our method, templates affect generation only through the conditional feature pathway, while the reverse parameterization follows DiffAb.

**Reverse process for amino acid types.** For residue types, DiffAb parameterizes the reverse categorical transition as:

$$p(s_j^{t-1} \mid \mathbf{R}^t, \mathcal{C}) = \text{Multinomial}\Big(F_\theta(\mathbf{R}^t, \mathcal{C})[j]\Big), \tag{S7}$$

where $F_\theta(\cdot)[j] \in \Delta^{19}$ outputs a categorical distribution over 20 residue types.

**Reverse process for coordinates.** Using the standard diffusion parameterization, DiffAb defines

$$p(\mathbf{x}_j^{t-1} \mid \mathbf{R}^t, \mathcal{C}) = \mathcal{N}\big(\boldsymbol{\mu}_\theta(\mathbf{R}^t, \mathcal{C})[j], \beta_t^{\text{pos}}\mathbf{I}\big). \tag{S8}$$

The mean is computed from a predicted positional noise vector $G_\theta(\mathbf{R}^t, \mathcal{C})[j]$ as

$$\boldsymbol{\mu}_\theta(\mathbf{R}^t, \mathcal{C})[j] = \frac{1}{\sqrt{\alpha_t^{\text{pos}}}}\left(\mathbf{x}_j^t - \frac{\beta_t^{\text{pos}}}{\sqrt{1 - \bar{\alpha}_t^{\text{pos}}}}\,G_\theta(\mathbf{R}^t, \mathcal{C})[j]\right), \quad \alpha_t^{\text{pos}} = 1 - \beta_t^{\text{pos}}. \tag{S9}$$

**Reverse process for orientations.** For orientations, DiffAb uses an isotropic Gaussian on $SO(3)$:

$$p(\mathbf{O}_j^{t-1} \mid \mathbf{R}^t, \mathcal{C}) = \mathrm{IGSO}(3)\Big(H_\theta(\mathbf{R}^t, \mathcal{C})[j],\ \beta_t^{\text{ori}}\Big), \tag{S10}$$

where $H_\theta(\cdot)[j] \in SO(3)$ predicts the denoised mean rotation at step $t - 1$.

**Equivariance (high-level).** The denoiser is $SE(3)$-equivariant: if the entire complex is rotated/translated, the predicted coordinates and orientations transform accordingly, while the residue-type distribution remains invariant—an essential property for molecular generation.

### S2.4 Training Objective

At training time, DiffAb samples a diffusion step $t \sim \text{Uniform}(\{1, \ldots, T\})$, corrupts the data state $\mathbf{R}^0$ into a noisy state $\mathbf{R}^t$ using the forward processes, and optimizes three terms (type / position / orientation). The dependence on $t$ is implicit through $\mathbf{R}^t$ and the noise schedules $\beta_t^\star$.

**Amino-acid type loss.** DiffAb minimizes the expected divergence $D_{\text{KL}}(q\|p)$ between the closed-form posterior $q(s_j^{t-1} \mid s_j^t, s_j^0)$ induced by the multinomial forward process and the model reverse kernel:

$$L_t^{\text{type}} = \mathbb{E}\left[\frac{1}{m}\sum_{j \in \mathcal{C}_{\text{cdr}}} D_{\text{KL}}\Big(q(s_j^{t-1} \mid s_j^t, s_j^0)\ \|\ p(s_j^{t-1} \mid \mathbf{R}^t, \mathcal{C})\Big)\right], \tag{S11}$$

where $m = |\mathcal{C}_{\text{cdr}}|$ is the number of generated residues.

**Position loss.** Let $\boldsymbol{\epsilon}_j \sim \mathcal{N}(\mathbf{0}, \mathbf{I})$ be defined through the reparameterization $\mathbf{x}_j^t = \sqrt{\bar{\alpha}_t^{\text{pos}}}\mathbf{x}_j^0 + \sqrt{1 - \bar{\alpha}_t^{\text{pos}}}\boldsymbol{\epsilon}_j$:

$$L_t^{\text{pos}} = \mathbb{E}\left[\frac{1}{m}\sum_{j\in\mathcal{C}_{\text{cdr}}}\left\|\boldsymbol{\epsilon}_j - G_\theta(\mathbf{R}^t,\mathcal{C})[j]\right\|_2^2\right]. \tag{S12}$$

**Orientation loss.** Let $\widehat{\mathbf{O}}_j^{t-1} = H_\theta(\mathbf{R}^t,\mathcal{C})[j]$ denote the predicted denoised rotation estimate. DiffAb measures the discrepancy by penalizing the deviation of the relative rotation $(\mathbf{O}_j^0)^\top\widehat{\mathbf{O}}_j^{t-1}$ from identity:

$$L_t^{\text{ori}} = \mathbb{E}\left[\frac{1}{m}\sum_{j\in\mathcal{C}_{\text{cdr}}}\left\|(\mathbf{O}_j^0)^\top\widehat{\mathbf{O}}_j^{t-1} - \mathbf{I}\right\|_F^2\right]. \tag{S13}$$

**Overall objective.** The total objective averages the three terms over a uniformly sampled diffusion step:

$$L = \mathbb{E}_{t\sim\text{Uniform}(1,\dots,T)}\left[L_t^{\text{type}} + L_t^{\text{pos}} + L_t^{\text{ori}}\right]. \tag{S14}$$

#### S2.5 Relation to Our Method

Our method uses DiffAb as the diffusion backbone, adopting the same diffusion variables, noise schedules, reverse parameterizations, and denoising losses; we do *not* modify the diffusion formulation itself. The key difference lies in conditioning: beyond the antigen-framework context, we optionally condition on candidate CDR templates $\mathcal{T}$ whose usefulness can vary. To make conditioning robust, we introduce a template-wise gating mechanism that suppresses uninformative template signals and prevents brittle dependence on them, while allowing the model to benefit from construction-positive or informative templates when available.

### S3 Implementation Details

#### S3.1 Model Details

Our implementation builds upon DiffAb and augments its conditioning with a template-conditioning module: we encode a pool of $K$ CDR-only *sequence* templates into continuous features, apply a template-wise gate to suppress uninformative or mismatched candidate signals, and inject the gated template features via multi-template cross-attention. We otherwise follow DiffAb for the single-residue and pairwise embeddings.

Non-CDR positions are padded with `[PAD]` for residue alignment, so the template stream provides only candidate CDR sequence information as an optional weak condition regulated by the learnable gate. We use dimensions 128 for both single-residue and template embeddings, 64 for pairwise embeddings, and $T = 100$ diffusion steps.

The template encoder $\mathcal{E}_\Theta$ is a learned amino-acid embedding (with `[PAD]`) followed by a 2-layer Transformer encoder (4 heads) and a linear projection, producing residue-aligned template features $\mathbf{Z}^{(k)} \in \mathbb{R}^{L\times d_t}$. The gate $g_\theta$ outputs one logit $\ell^{(k)}$ per template by applying an MLP to the feature-wise concatenation $[\mathbf{H}_t;\mathbf{Z}^{(k)}]$ and mean-pooling over residues: MLP$(2d \to 64 \to 64)$ with ReLU, followed by a linear head $64 \to 1$. We use a sigmoid-based STE to obtain $\tilde{\alpha}^{(k)} \in \{0,1\}$ in the forward pass. For fusion, we use a single post-norm cross-attention block (4 heads, dropout 0.1) with a 2-layer FFN (expansion ratio 4, GELU, dropout 0.1), and aggregate template messages by the gated normalized average as described in the main paper.

#### S3.2 Training Details

Training is performed in two stages. We first train the model on CDR-H3 only for 90k iterations with learning rate $1\times10^{-4}$, since CDR-H3 is the most structurally diverse and challenging region. Starting from this checkpoint, we fine-tune on the full (H1/H2/H3) dataset for an additional 300k iterations with a reduced learning rate of $5\times10^{-5}$.

We use Adam with $\beta_1 = 0.9$, $\beta_2 = 0.999$, batch size 16, and gradient clipping with maximum norm 100. Loss weights are set to $\lambda_{\text{rot}} = \lambda_{\text{pos}} = \lambda_{\text{gate}} = 1.0$ and $\lambda_{\text{seq}} = 5.0$; the gate loss is applied only to the gating module (with gradients stopped to the diffusion backbone). All experiments are conducted on a single NVIDIA A6000 GPU (48 GB). In total, we train for 390k iterations (about 94 hours). Unless specified, we use $K = 8$ candidate CDR templates. During training (**Trn-Mix**), we construct a mixed-quality pool per sample: (i) include one native template $S^\star$, (ii) add $\lceil\rho K\rceil$ homologous templates by mutating $S^\star$, and (iii) fill the remaining $K-\lceil\rho K\rceil-1$ slots with random templates. Here $\rho$ denotes the *fraction of homologous templates* in the $K$-template pool, and we use $\rho = 0.2$ unless otherwise specified.

For homologous construction, we mutate only CDR residues: each CDR position is independently mutated with probability $p_{\text{mut}}$ (we use $p_{\text{mut}} = 0.2$), and the substituted amino acid is sampled uniformly from the 19 alternatives (excluding the original residue). Random templates are formed by uniformly sampling amino-acid tokens at CDR positions and padding non-CDR positions with `[PAD]`. To encourage robustness, with probability $p_{\text{rand}} = 0.2$ we use the **all-random** setting (all $K$ templates are random); otherwise we use the above mixed-quality pool containing $S^\star$. At inference time, we apply a hard gate threshold $\tau = 0.4$ unless otherwise specified.

#### S3.3 Sampling Details

We set a fixed global seed (`seed_all(2025)`) once at the beginning of evaluation and do not re-seed thereafter. For each input, we call `model.sample` $N = 100$ times under identical conditioning and hyperparameters; the 100 samples differ only due to diffusion noise drawn sequentially from the same RNG stream (thus fully reproducible). We report the mean over all 100 samples, with no best-of-$N$ selection or post-hoc filtering: $\overline{M} = \frac{1}{100}\sum_{i=1}^{100} M(\hat{y}_i)$.

### S4 Train-Set Retrieval Protocol

To evaluate whether the learned conditioning behavior transfers beyond synthetic template pools, we construct a train-set retrieval setting using MMseqs2 candidates. For each test case, we use the heavy-chain non-H3 context as

TABLE S1: Train-set retrieval evaluation under the CDR-H3-only protocol. AAR is reported in percent and RMSD in Å.

| Train | Test | Gate | AAR↑ | RMSD↓ |
|---|---|---|---|---|
| RealBank | RealBank | OFF | 30.08 | 3.18 |
| RealBank | RealBank | ON | 31.76 | 3.13 |
| Trn-Mix | RealBank | ON | **39.50** | **2.76** |

TABLE S2: Sensitivity to the gate threshold $\tau$ under the CDR-H3-only protocol.

| $\tau$ | AAR (%,↑) |
|---|---|
| 0.2 | 36.78 |
| 0.3 | 38.30 |
| 0.4 | 39.47 |
| 0.5 | 39.01 |
| 0.6 | 38.33 |

the query and retrieve candidate templates only from the training set. The target CDR-H3 sequence and structure are not used as retrieval queries.

Retrieved candidates are filtered using the following criteria: sequence identity pident $\geq 80$, matched H3 length, deduplication, and top-4 ranking after filtering. This produces a hard-filtered real-bank candidate set. We evaluate three settings: (i) **Trn-RealBank / Tst-RealBank / Gate OFF**, (ii) **Trn-RealBank / Tst-RealBank / Gate ON**, and (iii) **Trn-Mix / Tst-RealBank / Gate ON**. The first two models are retrained on real-bank candidates to isolate the effect of the gate under hard-filtered retrieval conditions, while the third setting directly evaluates the original Trn-Mix model on train-set retrieval candidates.

This evaluation is more retrieval-based than the synthetic Tst-Rand and Tst-Oracle settings, but it should not be interpreted as a full end-to-end retrieval benchmark over a large external antibody database. Its purpose is to test whether the conditioning policy learned under mixed-quality template training remains effective when candidate templates are obtained from a train-set retrieval procedure.

### S5 Gate Threshold Sensitivity

We evaluate the sensitivity of NT-ABDiff to the inference-time hard gate threshold $\tau$ under the CDR-H3-only protocol. The main paper uses $\tau = 0.4$ as the default value. As shown in Table S2, AAR is stable around $\tau = 0.4$–$0.5$, supporting the use of $\tau = 0.4$ in the main experiments.

### S6 Seed-Level Variability

Table S3 reports five-seed variability for NT-ABDiff under the joint H1/H2/H3 protocol. The main paper reports benchmark means for all methods. Baseline standard deviations are not included because the reproduced benchmark logs provide single benchmark means for the compared baselines.

TABLE S3: Five-seed variability of NT-ABDiff under the joint H1/H2/H3 protocol. Values are reported as mean ± standard deviation.

| Setting | Metric | H1 | H2 | H3 |
|---|---|---|---|---|
| Rand | AAR (%) | 69.63±1.20 | 55.34±3.70 | 38.96±1.50 |
| Rand | RMSD (Å) | 1.141±0.10 | 1.188±0.10 | 3.065±0.20 |
| Rand | IMPROVE% | 49.00±3.40 | 31.11±4.60 | 25.52±2.80 |
| Oracle$^{\dagger}$ | AAR (%) | 70.39±0.90 | 54.69±4.30 | 39.42±1.20 |
| Oracle$^{\dagger}$ | RMSD (Å) | 1.121±0.10 | 1.122±0.30 | 2.994±0.10 |
| Oracle$^{\dagger}$ | IMPROVE% | 50.63±2.40 | 35.60±3.30 | 28.10±3.00 |

### S7 Scope of Template Perturbations and Gate Scores

The random templates used in the controlled experiments should be interpreted as extreme uninformative controls and as conditional perturbations during training. They are not intended to fully model the distribution of retrieval errors in external antibody databases. Instead, they provide a controlled way to test whether the denoiser can avoid depending on template signals when the provided candidates are uninformative.

Similarly, the gate score $\alpha$ should be interpreted as a construction-specific, context-conditioned template-usefulness score. It is not a calibrated uncertainty probability and is not trained from general biological reliability annotations. The gate is supervised using the mixed-quality construction labels, where native and homologous-like point-mutated candidates are labeled as construction-positive and random candidates are labeled as construction-negative. This supervision encourages the model to suppress uninformative template signals while allowing informative candidates to contribute through the template-conditioning pathway.

This work evaluates the conditioning module on a DiffAb-style sequence–structure diffusion backbone. Although the module is lightweight and modular, transfer to other antibody generative backbones is not evaluated here.

### S8 Qualitative Visualization

Fig. S1 provides qualitative examples of generated CDR-H3 conformations at the antigen interface. Each triplet shows the reference antibody and two generated samples. The CDR-H3 region is shown as orange sticks and the antigen interface is shown as blue sticks.

The reported $\Delta G$ values are computed using the same Rosetta-based scoring protocol used for the IMPROVE% metric in the main paper. These values should be interpreted as computational scores for qualitative comparison rather than experimentally validated binding affinities. The visualization is intended to illustrate interface-localized CDR-H3 conformations and should not be used as a substitute for quantitative evaluation or wet-lab validation.

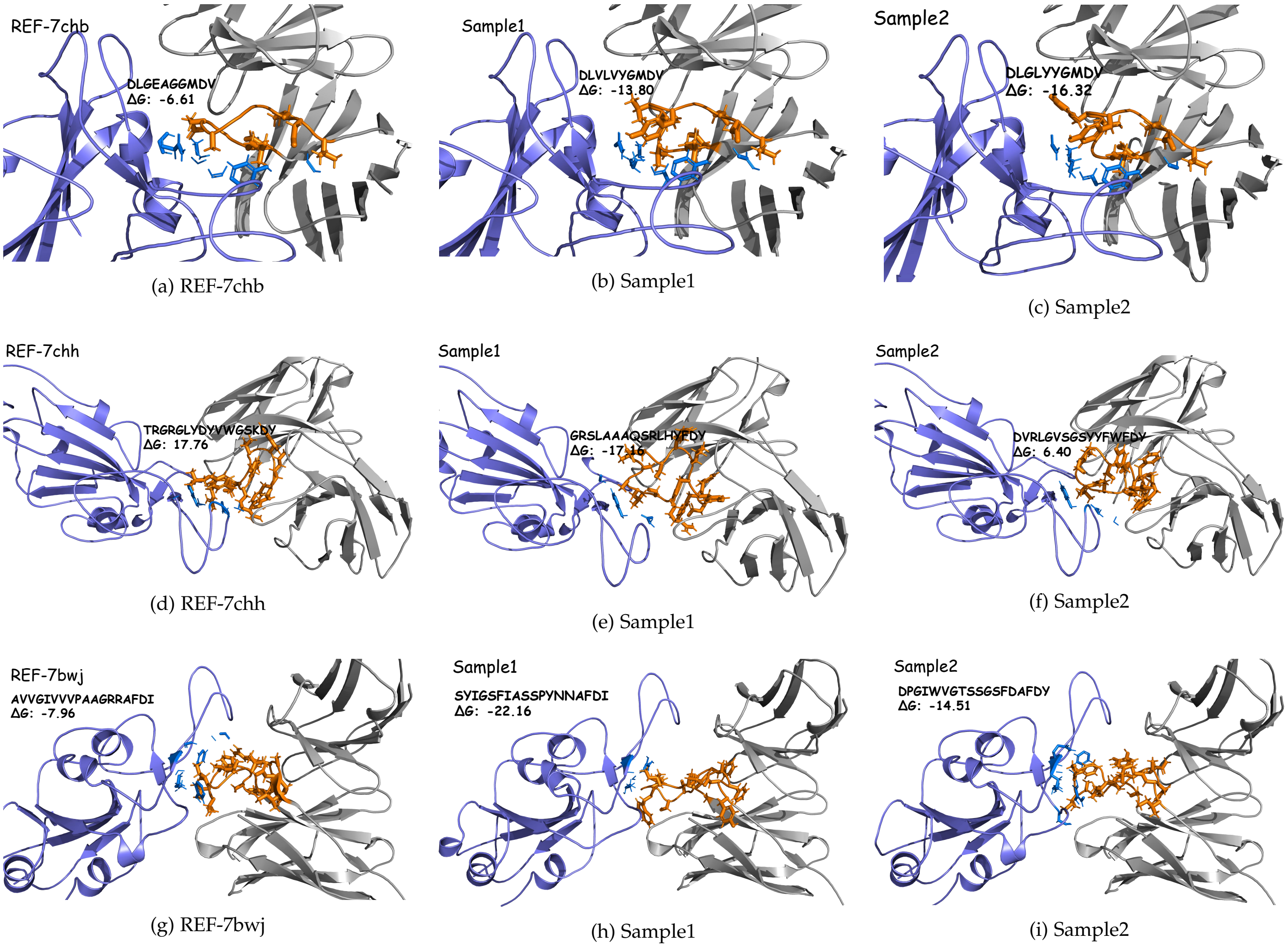


(a) REF-7chb (b) Sample1 (c) Sample2

(d) REF-7chh (e) Sample1 (f) Sample2

(g) REF-7bwj (h) Sample1 (i) Sample2

Fig. S1: Qualitative visualization of CDR-H3 conformations at the antigen interface. The CDR-H3 region is shown as orange sticks and the antigen interface is shown as blue sticks. Reported $\Delta G$ values are Rosetta-based computational scores and are not experimentally validated binding affinities.
